# Characterization of *Mycobacterium tuberculosis* rhodanese-like sulfurtransferase SseA and its newly identified activator Rv3284: A new pathway to exploit as a drug target?

**DOI:** 10.1101/2025.02.04.636439

**Authors:** Giulia Di Napoli, Alex Fissore, Edoardo Salladini, Eleonora Raccuia, Simonetta Oliaro-Bosso, Alessia Ruggiero, Rita Berisio, Milagros Medina, Adrian Velazquez-Campoy, Salvatore Adinolfi, Mauro Marengo

**Affiliations:** Department of Drug Science and Technology, University of Turin, Turin, Italy; Institute of Biostructures and Bioimaging, CNR, Naples, Italy; Department of Biochemistry and Molecular and Cellular Biology, School of Sciences, and Institute for Biocomputation and Physics of Complex Systems, University of Zaragoza, Zaragoza, Spain; Instituto de Investigación Sanitaria Aragón (IIS Aragón), Zaragoza, Spain; Centro de Investigación Biomédica en Red en el Área Temática de Enfermedades Hepáticas y Digestivas (CIBERehd), Madrid, Spain

**Keywords:** SseA, SufE, *Mycobacterium tuberculosis*, Enzyme, Thiosulfate-sulfurtransferase

## Abstract

Tuberculosis (TB) remains a critical global health challenge, with *Mycobacterium tuberculosis* (Mtb) causing 10.8 million cases and 1.25 million deaths in 2023. Key issues include the dormant state of Mtb, resistant to drugs and immune responses, and the emergence of multi-drug-resistant strains. This underscores the need for new therapeutic targets and deeper research into Mtb pathogenesis and immunology.

A potential drug target is the enzyme thiosulfate-sulfurtransferase SseA, which plays a role in macrophage infection by Mtb and its resistance to oxidative stress. SseA belongs to the rhodanese-like enzyme family, which catalyzes sulfur transfer reactions essential for Mtb survival.

In our research, we identified a new protein (Rv3284), hereinafter referred to as SufE_Mtb_ due to its high homology with *E. coli* SufE, that interacts with SseA and modulates its activity. Sequence analysis and AI molecular modelling revealed detailed insights into their interaction that can contribute to the modulation of SseA activity. This research provides a mechanistic explanation to the need of a partner for SseA activation. Indeed, we propose that SufE_Mtb_ enhances SseA enzymatic function by binding to its non-catalytic N-terminal domain and bringing the active sites of the two proteins in close proximity, thus preparing for the activation-enhancing conformational change in a regulatory loop of SseA. This interaction is crucial for the effective enzyme activity and the maintenance of redox homeostasis in Mtb, making the SseA-SufE_Mtb_ protein complex a potential target for new TB therapies.

## Introduction

Tuberculosis (TB) remains a major global health issue and one of the leading causes of death worldwide. The etiologic agent, *Mycobacterium tuberculosis* (Mtb), caused 10.8 million cases of TB and 1.25 million deaths in 2023 (WHO). The most urgent problems of TB infections are the capacity of Mtb to enter a dormant non-replicating form through complex modifications of its cell wall [1], which is resistant to drugs and immunoresponse. Also, Mtb has an extraordinary ability to develop multi-drug resistant (MDR) and extensively drug-resistant (XDR) forms [2]. These events have further emphasized the need to discover new essential functions, in pathways different from those targeted by conventional antibiotics to combat this disease [2]. This goal requires more basic research to better understand Mtb pathogenesis and immunology, and to identify new targets for diagnostics, drugs and vaccines [3].

Although its physiological role in Mtb has not been clarified yet, the putative thiosulfate-sulfurtransferase SseA appears as a potential drug target candidate since it has been demonstrated to be involved in Mtb macrophage infection [4] and is overexpressed in MDR and XDR Mtb strains [5]. It is suggested to play a fundamental role in the molecular pathways associated to oxidative stress that are necessary for Mtb to resist oxidation during infection and for the bacterial cytosolic thiol homeostasis [6]. SseA belongs to a large family of rhodanese-like enzymes that form a transient sulfane sulfur during catalysis and are able to use a number of substrates, including low-molecular-weight thiols [6]. In particular, two types of sulfurtransferases with tandem rhodanese domains have been described. The first group, widely distributed in both prokaryotes and eukaryotes [7–9], referred to as thiosulfate:cyanide sulfurtransferases (TST), showed *in vitro* specificity for thiosulfate to produce thiocyanide [7], whereas the second group, referred to as 3-mercaptopyruvate sulfurtransferases (MST) uses 3-mercaptopyruvate as sulfur donor to cyanide to obtain pyruvate and thiocyanate as final products [10,11]. As for SseA, the protein shows the TST characteristic motif CRXGX[R/T][10].

According to the accepted mechanism [12], TST enzymes facilitate the transfer of sulfur from thiosulfate (donor) to cyanide (thiophilic acceptor) through a double displacement reaction. Initially, TST/rhodanese acquires a sulfane sulfur atom from a donor (e.g., thiosulfate), leading to the creation of a covalent enzyme-sulfur intermediate (E-S) characterized by a persulfide bond at the sulfhydryl group of the reactive cysteine in the active site. Subsequently, the persulfide sulfur is transferred from the enzyme to the cyanide, restoring the enzyme to its original form [9,13,14].

There is strong evidence that TST and MST are evolutionarily related, as they are able to interact with the same substrate (cyanide, thiosulfate, 3-mercaptopyruvate), albeit with distinct kinetics and different affinities, and show a striking similarity in amino acid sequences around the active site (66% of sequence homology between TST and MST in rat liver) [15].

Among rhodanese-like proteins, a thorough structural and enzymatic characterization was carried out on SseA from *E. coli* [16].

Next to the SseA gene (Rv3283), along the Mtb genome sequence there is a neighboring gene (Rv3284) that encodes for a yet uncharacterized protein. Commonly, proteins encoded by neighboring genes are expressed and work together in a specific metabolic pathway. It was thus tempting to make the working hypothesis that Rv3284 protein collaborated with SseA in cysteine desulfuration, by providing the sulfur atom during cyanide detoxification. In this study we report the purification and the biochemical characterization of the rhodanese-like protein SseA and of its neighbor gene product Rv3284, that showed high homology with *E. coli* SufE and is able to display a significant increase of SseA desulfurase activity. Biophysical characterization and computational analysis were instrumental to prove that SufE-like protein modulation is mediated by a direct binding to SseA with a 1:1 stoichiometry. Molecular modeling provided insights into a plausible activation mechanism of SseA induced by the interaction between the two proteins. Indeed, this interaction mode that provides contiguity between the two active sites and prepares SseA for the unlocking of a regulatory loop that limits the catalytic site access in SseA and is absent in the homologous *E. coli* protein.

## Results

### Identification of a putative SufE-like gene in Mtb

Through a bioinformatics search on the Mtb genome, we identified a neighbor sequence to the SseA coded as Mtb Rv3284. This sequence shares high homology (60%) with *E. coli* SufE, with virtually no insertions/deletions (**Fig. 1**). *E. coli* SufE is a sulfur transfer protein that promotes a significant increase in the catalytic activity of the cysteine desulfurase enzyme involved in transferring sulfur atoms to synthesize Fe-S cluster [25]. Different than observed in *E. coli*, Rv3284 is not part of the Suf operon, but it presents the well conserved Cys55 (corresponding to the Cys51 in *E. coli* SufE) essential for SufE to accept S_0_ via a thiol exchange mechanism [26–29]. Genomic neighborhood is indeed a reliable indicator for proteins to be involved in the same functional pathways. Consistently, the STRING database predicts the ability of SseA and Rv3284 to form a complex (data not shown). Due to its features, Rv3284 protein is here denominated as SufE_Mtb_.

**Figure 1:**
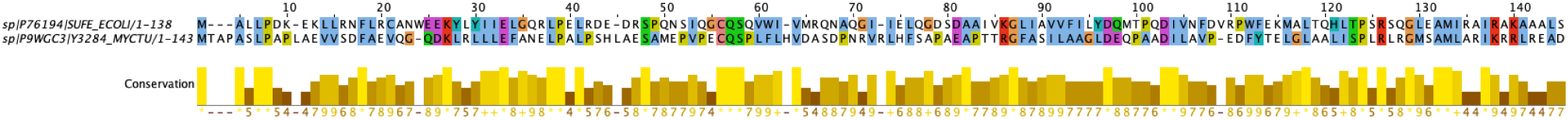
Sequence alignment of *E. coli* SufE and Rv3284 SufE_Mtb_ from *M. tuberculosis*. Color-coded is used to highlight similarities. All G and P residues are colored in orange and yellow, respectively. Other coloring is used to indicate a conserved property according to the following convention: blue, hydrophobic; red, positive; purple, negative; green, hydrophilic. The conservation is highlighted by the yellow bar below the sequences. The aminoacid conservation pattern is shown from light (most conserved) to dark yellow (least conserved).

### Purification of SseA and SufE_Mtb_

The setup of convenient purification protocols, leading to the production of soluble isolated highly pure proteins, is fundamental to addressing their individual properties, as well as their interaction properties. The overexpressed SseA and SufE_Mtb_ proteins were produced using pETM-20 and pET-24 plasmids, respectively. SseA and SufE_Mtb_ were purified from the bacterial cell extracts as fusion protein with His-tagged thioredoxin and with His-tagged glutathione-S-transferase, respectively, by making use of chromatographic approaches and taking advantage of the inserted six-histidine tag. The fusion proteins underwent digestion by using the His-tagged TEV protease, followed by an IMAC chromatography step that allowed to obtain soluble and pure tag-free SseA and SufE_Mtb_, as shown by SDS-PAGE (**Fig. 2A**).

**Figure 2:**
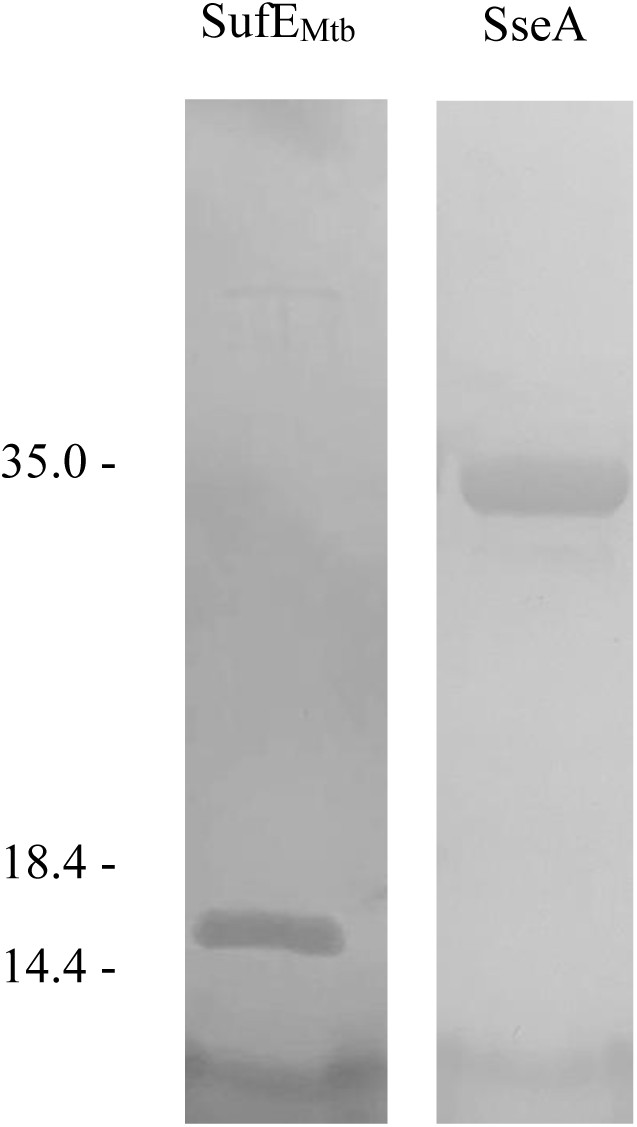
SDS-PAGE of recombinant proteins and Gel filtration. SDS-PAGE showing the purity of the proteins. The band at 14.4 kDa corresponds to SufE_Mtb_ purified to homogeneity, while the band at approximately 35 kDa corresponds to SseA purified to homogeneity.

The purified Mtb SseA was also analyzed by SEC, highlighting a single peak eluting at a volume compatible with the molecular weight of the monomer (33 kDa), as inferred from the calibration curve (data not shown).

### SseA sulfurtransferase activity

The purified SseA was used for enzymatic activity measurements in the presence of thiosulfate as a substrate, clearly highlighting the ability of the enzyme to transfer sulfur from the substrate to the cyanide acceptor molecule, most probably by the formation of a sulfur-substituted form of SseA (**Fig. 3A**). However, sulfur transfer from thiosulfate to cyanide appears moderate, calling for investigating and identifying a more appropriate substrate as a sulfur donor. In this frame, a number of potential acceptors (3-mercaptopyruvate, glutathione, cysteine) might be used instead of cyanide, which is not physiologically present in cells.

**Figure 3:**
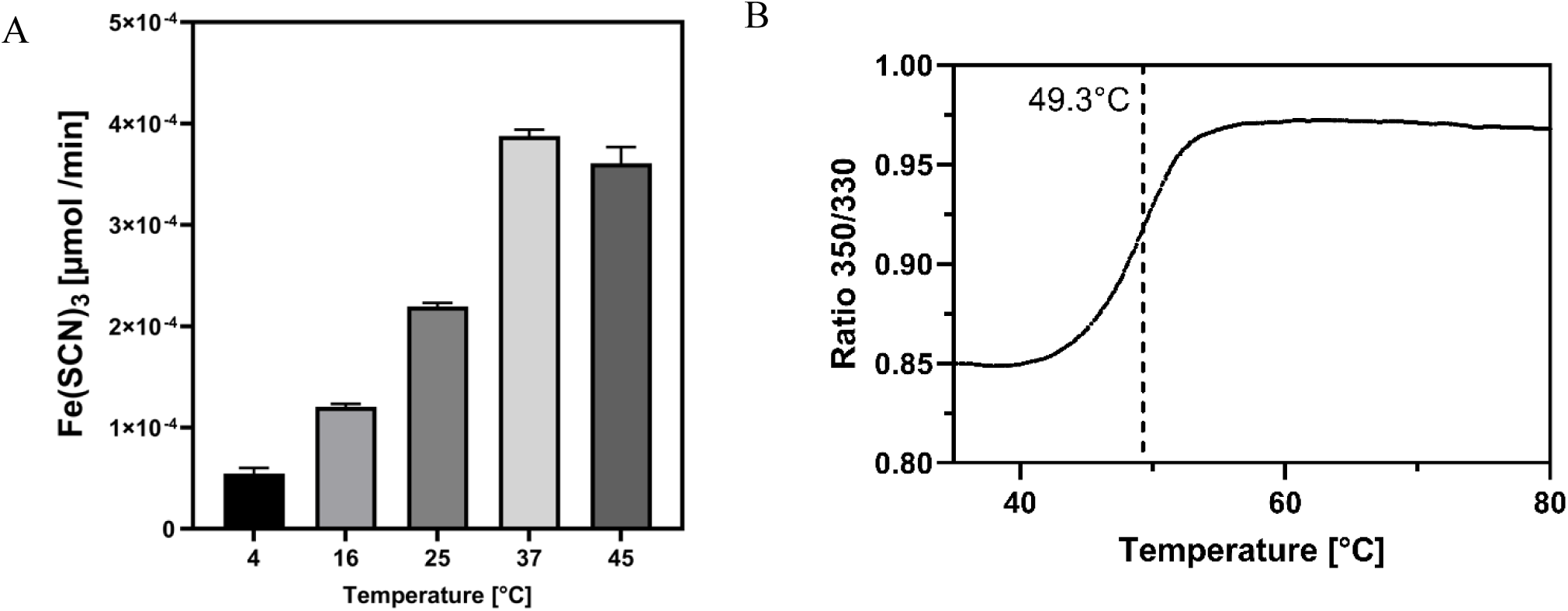
Temperature Dependence of Mtb SseA Enzymatic Activity and Protein Stability. Panel A, bar graph illustrating the increase of enzymatic activity (µmol of Fe(SCN)₃ produced per minute) up to 37 °C. Panel B, fluorescence trace analysis (350/330 nm) for protein thermal denaturation. Thermal melting curve was used to determine the melting temperature (T_m_), which was found to be 49.3 °C.

The optimal temperature for SseA enzymatic activity was identified by performing activity measurements over a range of temperatures. Results clearly show that the enzyme exhibits maximum catalytic activity at 37 °C, while a slight decrease is detected at 45 °C. No significant measurements could be performed at higher temperature, due to protein precipitation (**Fig. 3A**). These results are consistent with the enzyme melting temperature (T_m_) of 49.3 °C (**Fig. 3B**), as measured by Tycho (NanoTemper Technologies, Munich, Germany). The decrease in activity observed at temperatures above 37 °C most likely reflects the enzyme approaching its thermal threshold, where structural destabilization begins to occur.

### SseA activity is modulated by SufE_Mtb_

A potential effect of SufE_Mtb_ on SseA enzymatic activity could be predicted by their gene neighboring location, that appears as a reliable indicator for functional networks of proteins expressed and working together within a specific metabolic pathway. In this context, it has to be highlighted that SufE from *E. coli* promotes a considerable boost in the cysteine desulfurase activity of the iron-sulfur cluster biogenesis Suf machinery, contributes to transfer sulfur to the scaffold for the newly synthesized Fe-S cluster [30], and shares a high degree of sequence homology with SufE_Mtb_.

To investigate this interaction, SseA enzymatic activity was measured using thiosulfate as a substrate in the presence of increasing concentrations of SufE_Mtb_ (10–100 µM). The results revealed that SufE markedly enhances SseA sulfurtransferase activity, achieving a 4-fold increase at 100 µM SufE_Mtb_ (Fig. 4).

**Figure 4:**
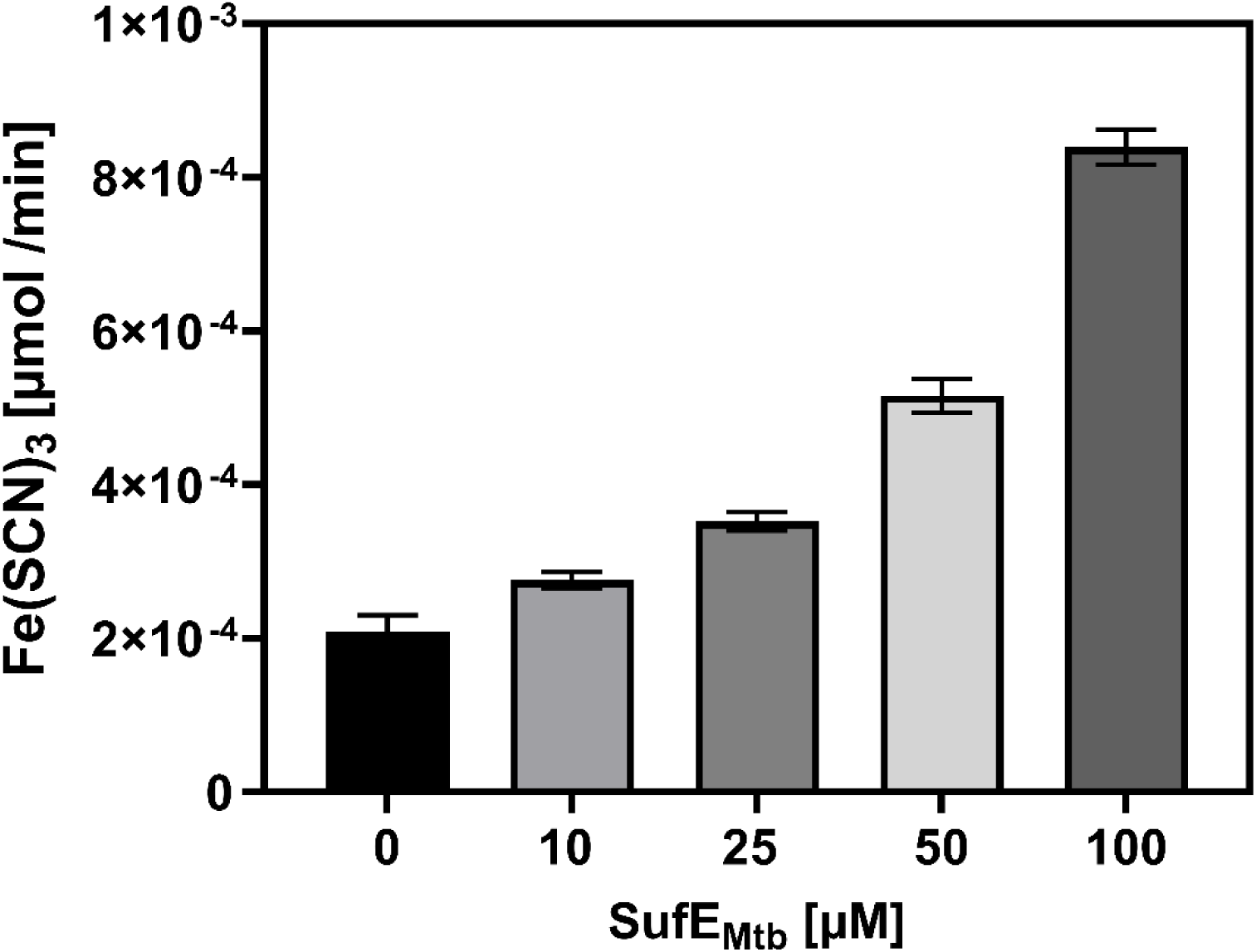
Modulation of Thiosulfate:Cyanide Sulfurtransferase Activity of *Mycobacterium tuberculosis* SseA by SufE_Mtb_. The thiosulfate:cyanide sulfurtransferase (TST) activity of purified Mtb SseA was assessed in the presence and in the absence of varying concentrations of SufE_Mtb_, at 20°C. The activity was measured by monitoring the conversion of thiosulfate and cyanide into SCN^−^ and subsequently revealed as Fe(SCN)_3_.

### SseA and SufE_Mtb_ complex formation

To address whether the activity increase resulted from a direct protein-protein interaction, MST measurements were carried out in the presence of 10 nM SseA labeled with the RED-NHS fluorescent dye and increasing concentrations (0.0015–50 μM) of the label-free SufE_Mtb_. Changes in the detected fluorescence signal during MST measurements provided experimental evidence of the protein-protein interaction, that appeared relevant to the biological activity of SseA and allowed to estimate a *K*_d_ value of 3.0 ± 0.7 μM for this interaction (**Fig. 5**).

**Figure 5:**
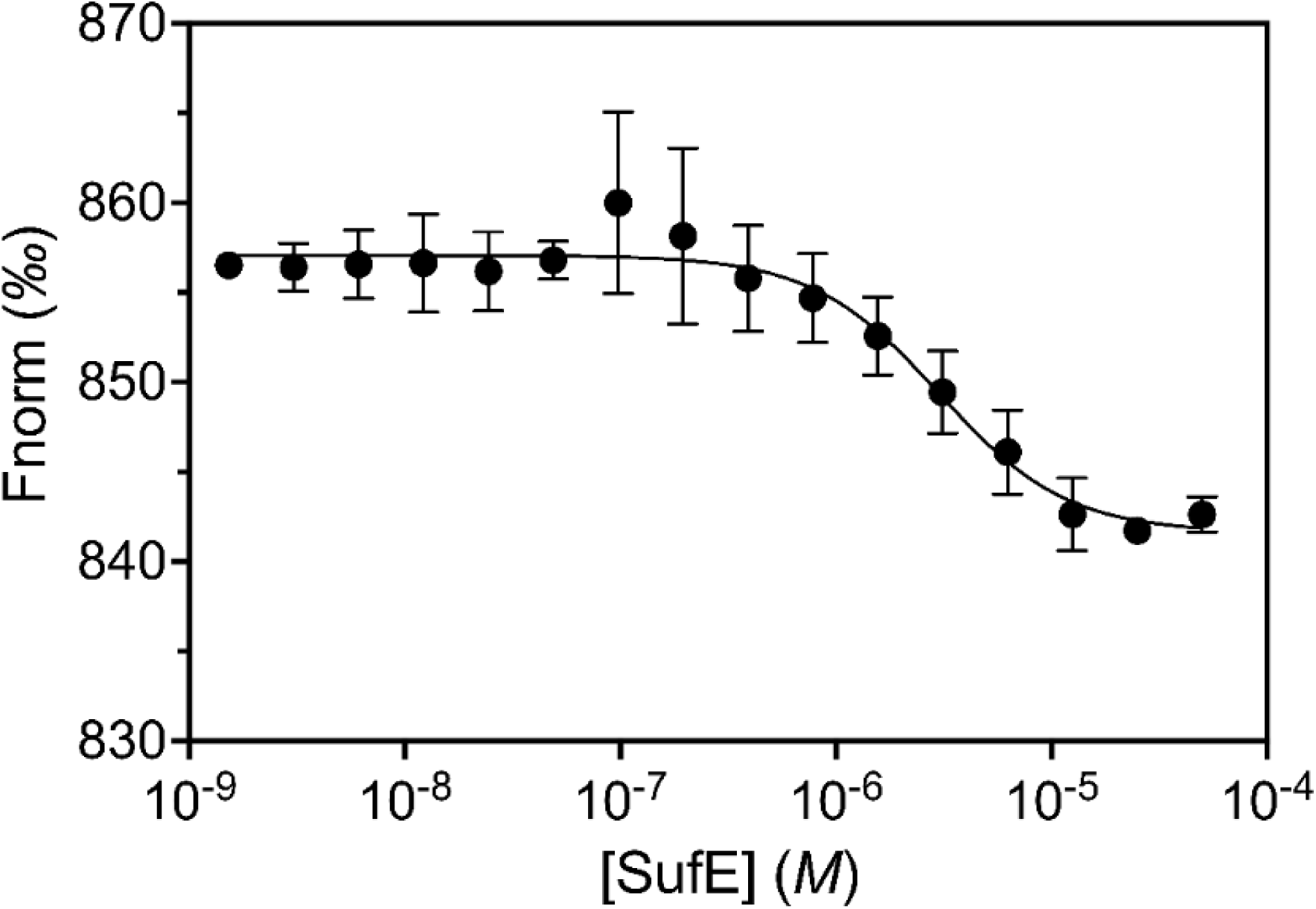
SseA-SufE_Mtb_ interaction via Microscale Thermophoresis (MST). Normalized fluorescence of labeled SseA when titrated with different amounts of unlabeled SufE_Mtb_. Measurements were performed at 25 °C using the NANO red LED at 100% excitation power and the MST power at 40%. The cold region was from −1.0 to 0 seconds and hot region from 4.0 to 5.0 seconds. Fitting curves were obtained by analysis with the MO Affinity Analysis v3.0.5 software.

The nature of the interaction between SseA and SufE was also investigated by isothermal titration calorimetry (ITC), that allows measuring binding constants with high accuracy. Titration of SseA with SufE_Mtb_ resulted in an exothermic binding, which could fit with a 1:1 binding stoichiometry model, and provided a *K*_d_ of 3.3 µM ± 0.6 μM (**Fig. 6**). These results are in agreement with the data obtained by MST measurements and confirm that the proteins interact with a moderate affinity (low µM range) through an enthalpically driven process (Δ*H* = –11.6 kcal/mol).

**Figure 6:**
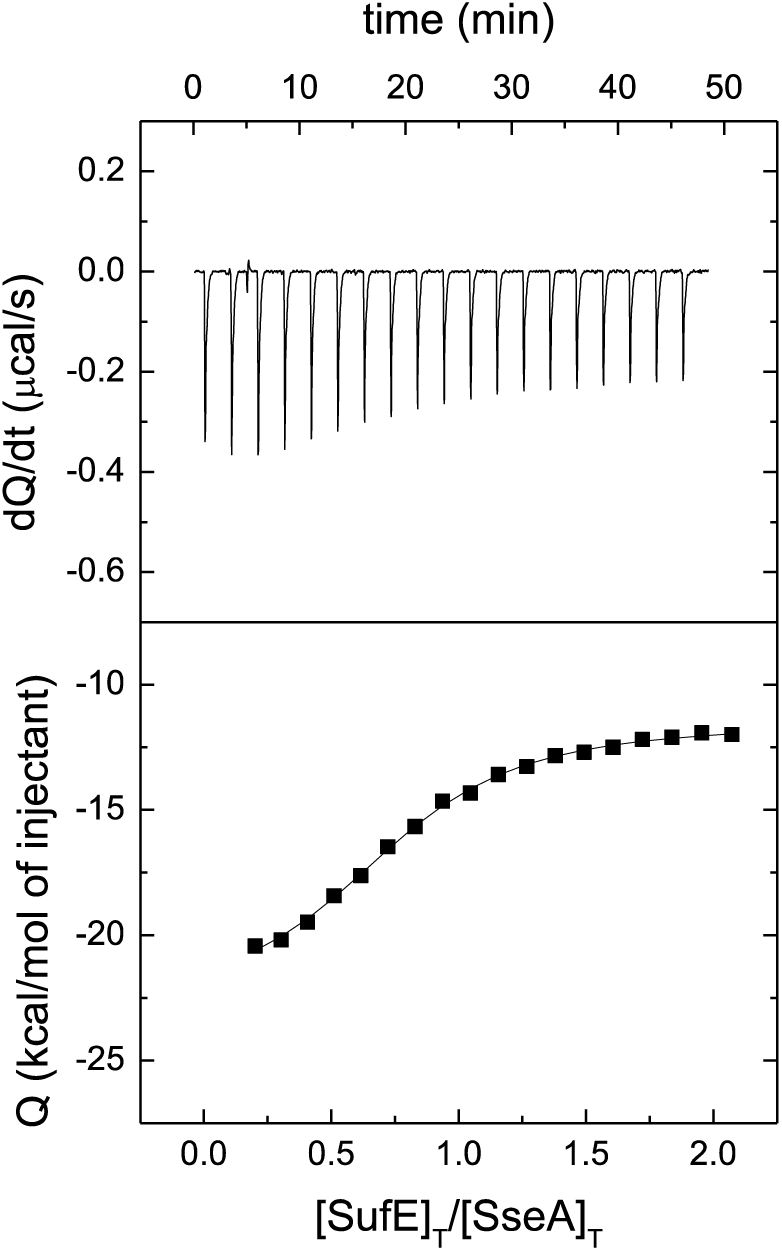
SseA binding to SufE_Mtb_ protein through calorimetric assays. Calorimetric titrations corresponding to the interaction of SseA with SufE_Mtb_ in phosphate buffer at 25°C. Upper panels show the thermograms (thermal power as a function of time) and lower panels show the binding isotherms (titrant-normalized injection heat effect as a function of the molar ratio within the calorimetric cell). Non-linear least-squares fitting (continuous lines in the lower panels) allowed estimation of the interaction parameters (dissociation constant, *K*_d_; observed interaction enthalpy, Δ*H*; and apparent stoichiometry, *n*).

The binding stoichiometry of 1:1 between SufE_Mtb_ and SseA, as determined by calorimetric assays, is indeed surprising given the structural complexity of SseA, which contains tandem N- and C-terminal rhodanese domains (Fig. 7A). To address these discrepancies, a sequence alignment of the two rhodanese domains of SseA was carried out in an attempt to elucidate the underlying mechanisms of interaction.

**Figure 7:**
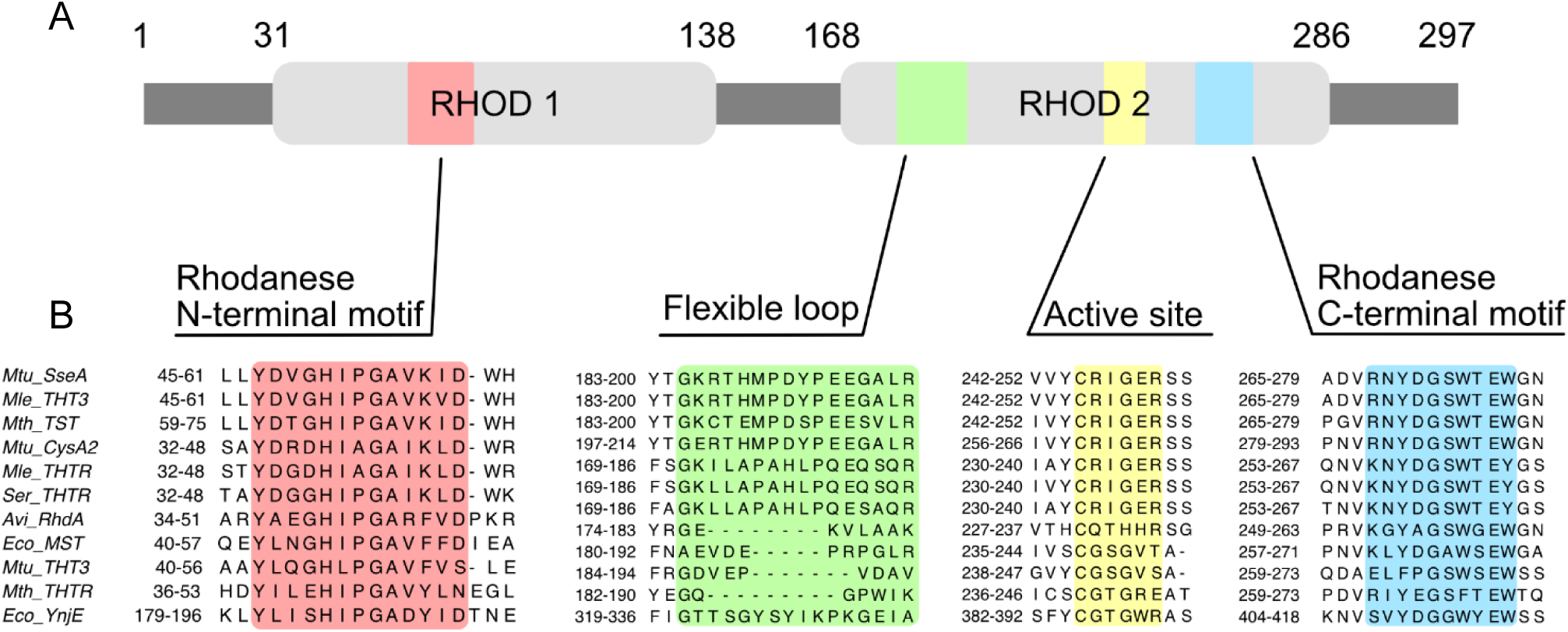
Sequence conservation of relevant regions of SseA rhodanese domains. (A) Domain architecture of SseA (A) and (B) sequence alignment of selected regions of SseA rhodanese domains with rhodanese-like proteins orthologues. These regions include N (red) and C (blue) terminal motifs of Rhod1 and Rhod2, the active site (yellow) and the flexible loop (green). Numbering in (A) corresponds to the *E. coli* protein.

The multiple alignment of rhodanese and rhodanese-like proteins, focusing on the two rhodanese domains and the active site [10] is shown in **Fig. 7**. This analysis reveals that, while sequence homology is maintained across the represented domains, certain sequences lack the same disordered loop. Specifically, sequences Avi_RgdA, Eco_MST, Mtu_THT3, and Mth_THTR exhibit gaps in the region highlighted in green. Considering the significance of the observed differences in the interactome of these proteins and the crucial role of the disordered region for the enzyme functionality highlighted in this study, a more detailed investigation into this region will be pursued in future research.

### The molecular Ssea:SufE_Mtb_ interacting complex

Mtb SseA folds in two domains with classical rhodanese-like fold. The N- and C-terminal domains are made, respectively, by residues 1-132 and 161–296 and connected by a linker (residues 133 –161) that wraps around the N-terminal domain and interacts with both domains (**Fig. 8A**)[31]. RMSD between the PDB and AF structures for SseA is as low as 0.5 Å, with major differences in the 185-201 loop. Consistently, this loop shows the highest thermal factors in the crystal structure and the lower confidence in prediction in the AF model (**Fig. 9A**). A cleft between the two rhodanese-like domains houses the proposed active site (**Fig. 8A, right**), with residues from both domains lining the catalytic Cys245 and the Ser251 assisting catalysis. On its side, the SufE_Mtb_ folds in an αβ compact architecture. The cysteine expected to be involved in persulfide-formation, Cys55, sits at the tip of an external loop with its sidechain buried from solvent exposure (**Fig. 8B**). Noticeably, Cys55 shows the lowest pLDDT confidence value within the AF model (87 for its Cα), whereas corresponding average values for the loop containing it and for the remaining protein are higher (respectively over 93 and 98) (**Fig. 9B**), in agreement with its expected displacement towards the surface during catalysis. The rest of the loop contains three Pro residues, which are not conserved in the bacterial *E. coli* structure[32]. Models for the SseA: SufE_Mtb_ interaction as produced by AF3 and AF2 were virtually identical (**Fig. 10A** and **10B**), and AF3 models showed predicted template modelling and interface predicted template modelling scores respectively of 0.83 and 0.69. These values are indicative of overall complex predicted fold being highly similar to the true structure as well as with a confident high-quality of the prediction. The model shows that SufE_Mtb_ interacts with the N-terminal rhodanese domain of SseA in a binding fashion that brings the loop of SufE_Mtb_ embedding Cys55 at the entrance of the SseA cleft between the N- and C-terminal domains (**Fig. 10C**). Specifically, Glu42 and Asp43 of SseA form hydrogen bonds with Arg124 of SufE_Mtb_. Another interaction is formed between SseA Asp43 and SufE_Mtb_ Thr83. Notably, Lys99 of SseA hydrogen bonds with Gln56 of SufE_Mtb_, adjacent to Cys55. The only region of the SseA C-terminal domain in contact with SufE_Mtb_ is the 185-201 loop (**Fig. 10C**). Its Tyr193 and Glu195 residues form hydrophobic and salt bridge interactions with Leu123 and Arg126, respectively, located on the α-helix following the reactive SufE_Mtb_ Loop. In this conformation, the SseA 185-201 loop prevents access of the SufE_Mtb_ reactive loop to its active site. However, the low pLDDT score of the SseA loop within the SseA:SufE_Mtb_ complex (average Cα values of 50) as compared to the free protein (average Cα of 75) suggests a high degree of flexibility upon protein-protein interaction is produced (**Fig. 9A** and **9C**). This suggests its position can highly vary to allow Cys55 of SufE_Mtb_ to approach the SseA active site. In fact, in SufE_Mtb_, the loop containing Cys55 and the portion of the helix in contact with SseA also show less confidence (average pLDDT Cα values below 74) than the rest of the structure (pLDDT Cα values over 98). This further agrees with the SufE_Mtb_ loop allowing flipping of Cys55 from buried towards solvent exposed to become accessible to the SseA active site. The presented model envisages these flexible loops within the complex to be more mobile than in the free proteins (**Fig. 9**), and, therefore, more prone to being displaced to achieve a catalytic competent organization for the transfer of sulfur from the active site of SseA to the thiol at Cys55 of SufE_Mtb_. Therefore, they appear potential early stage snapshots in the SseA:SufE_Mtb_ interaction for subsequent catalysis.

**Figure 8:**
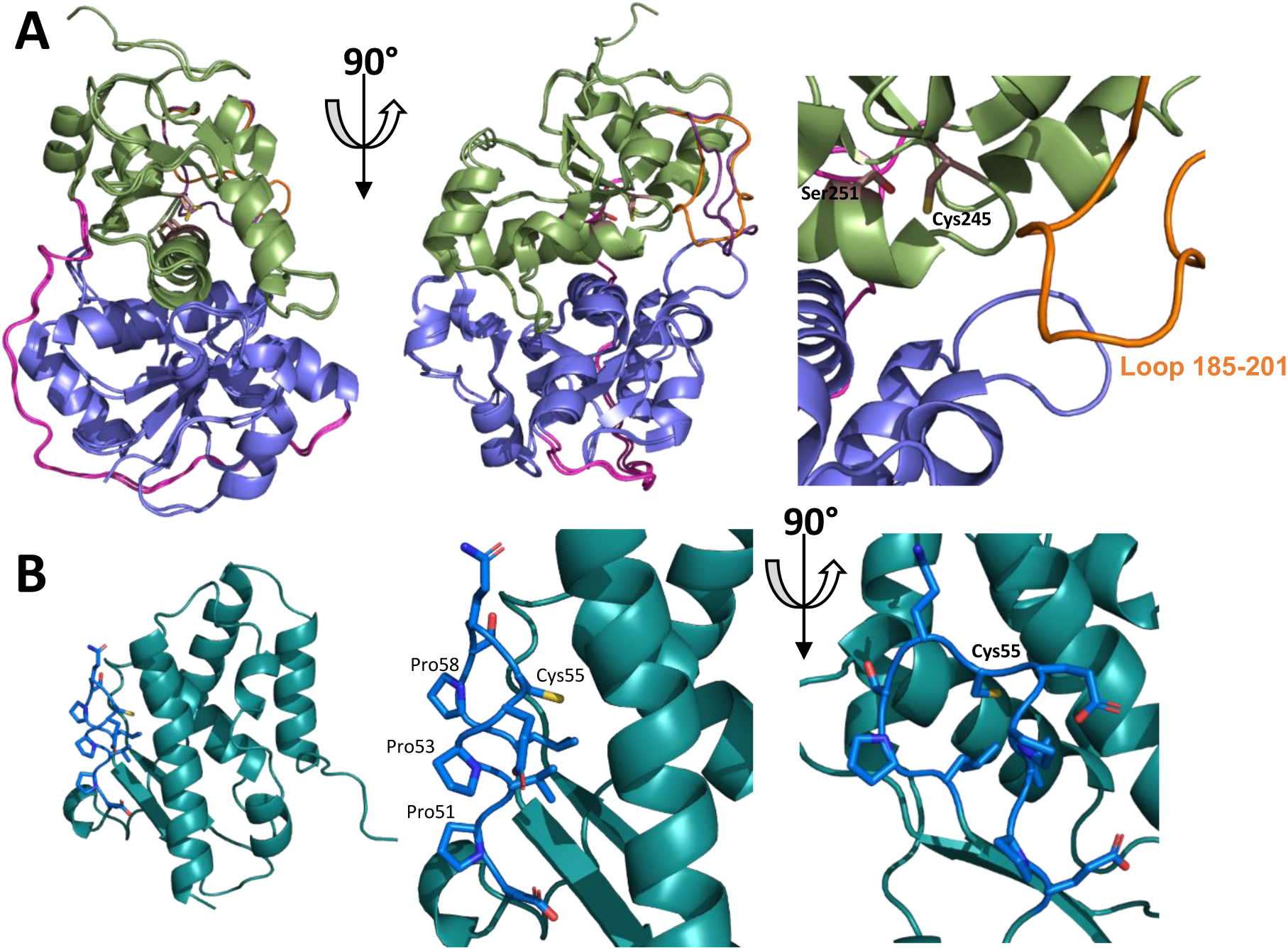
Structural models for SseA and SufE_Mtb_ from *M. tuberculosis*. (A) Overlapping of the crystallographic (3HZU) and AF (AF-P9WHF7-F1) structures for SseA (left panel) and detail of the active site organization (right panel). N- and C-terminal domains are respectively colored in lavender and green; their connecting loop is in magenta and the Loop 185-201 is colored in orange and violet respectively for the crystal and AF structures. Sidechains of Cys245 and Ser251 at the active site are shown in sticks with carbons in brown. (B) Overall AF structural model for SufE_Mtb_ (AF-P9WHF7-F1) (left panel) and detail of the loop (residues shown in sticks) containing the redox active Cys with its C atoms highlighted in dark blue, while the rest of the protein is in teal.

**Figure 9:**
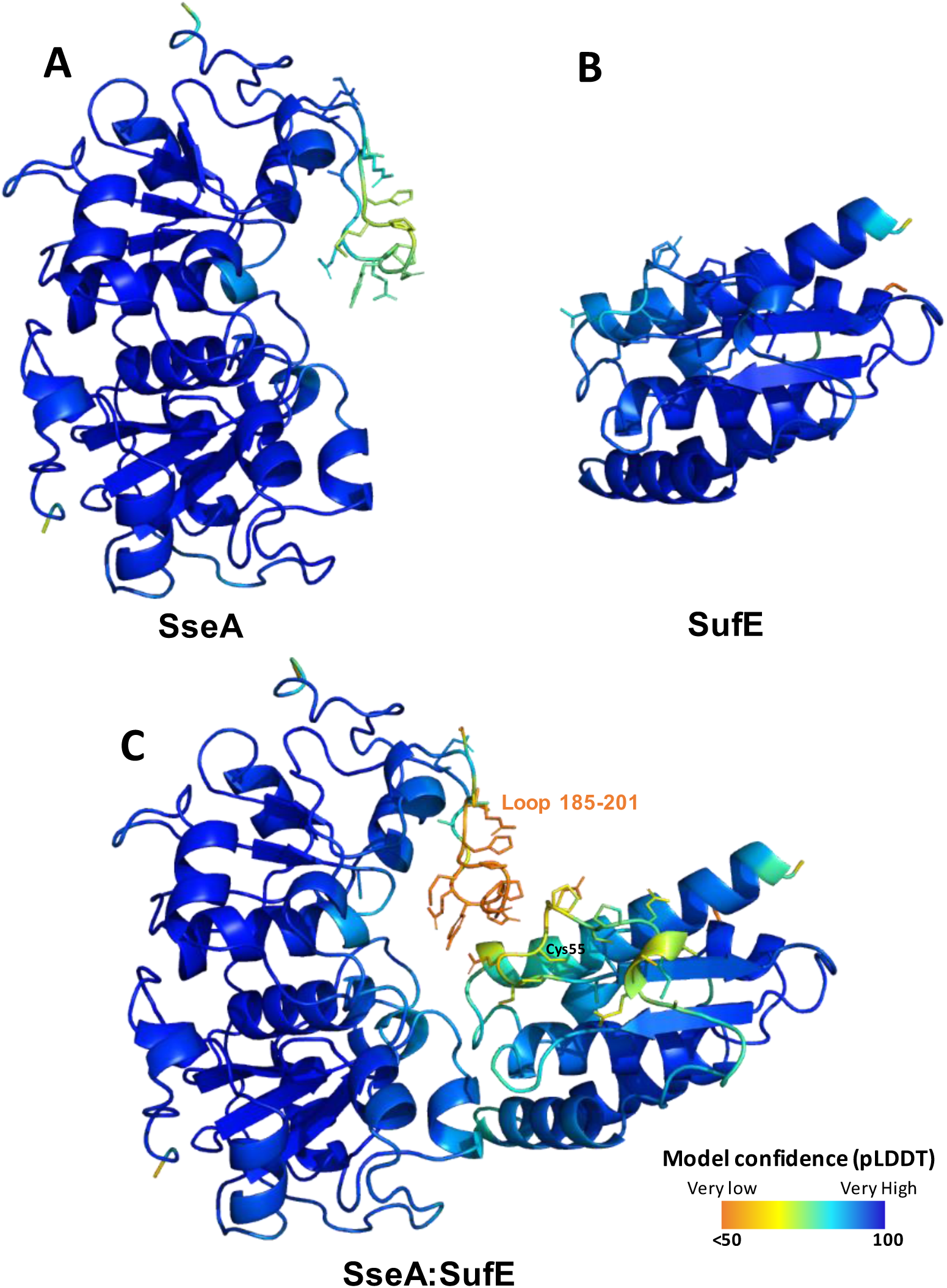
Residue model confidence of free proteins in comparison with the SseA:SufE_Mtb_ complex. **(A)** AFDB model for SseA (AF-P9WHF7-F1), (B) AFDB model for SufE_Mtb_ (AF-P9WHF7-F1) and (C) AF3 first model of the SseA:SufE_Mtb_ complex colored per-residue model confidence score (pLDDT) which ranges from 50 (orange) to 100 (dark blue). Side-chains of key loops are shown in sticks.

**Figure 10:**
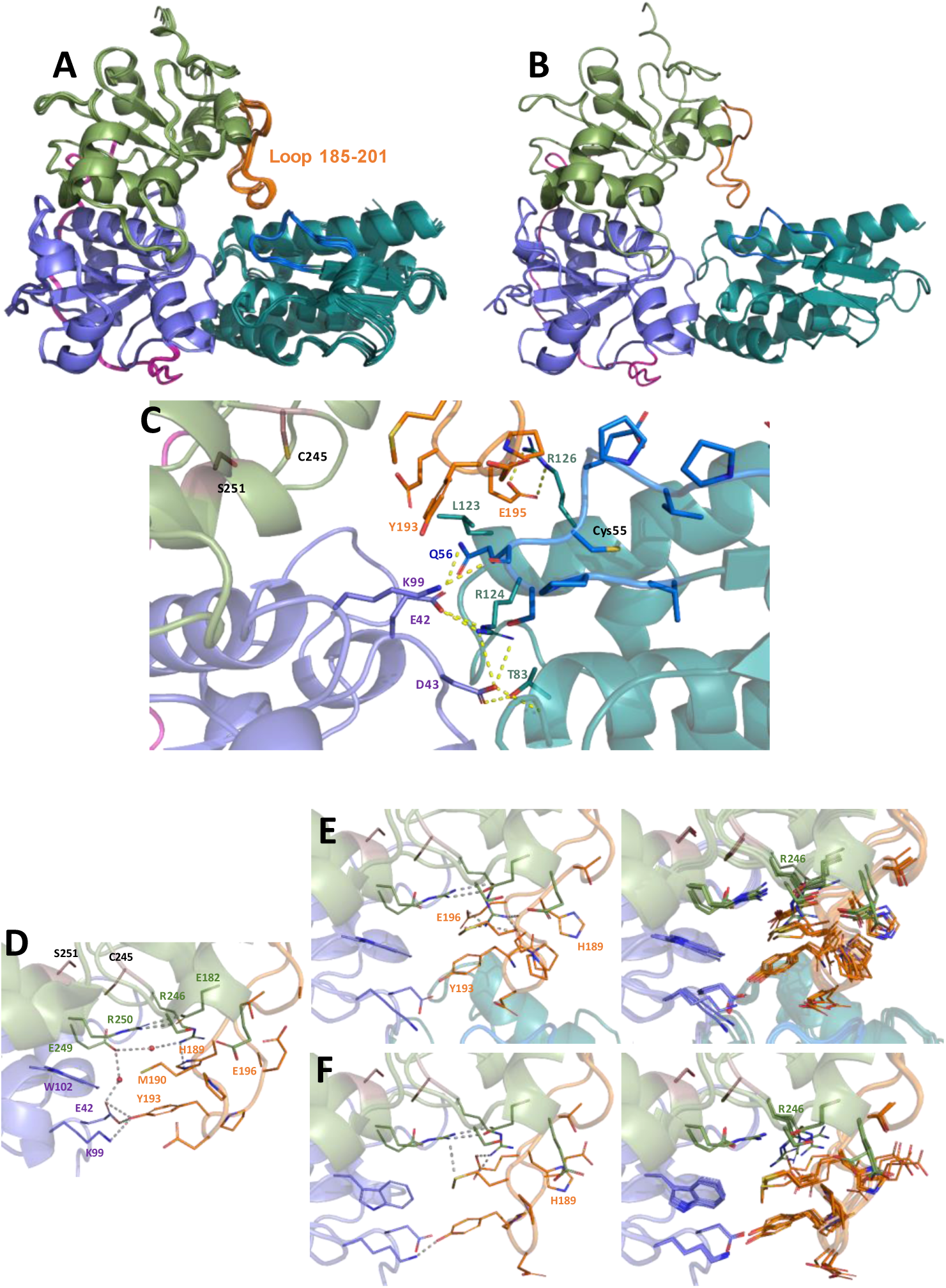
Structural model for the coupling of SseA and SufE_Mtb_ from *M. tuberculosis* under non-reactive conditions. (A) Overlapping of five replicates of the modelling as carried out by AF3. (B) Model produced by AF2 Multimer Colab. (C) Detail of the interface of the SseA:SufE_Mtb_ interaction in the first model provided by AF3. Detail of the conformation of contacts of the 185-201 loop of SseA with its core N- and C-terminal domains in: (D) The MtbSseA crystal structure (PDB 3ZHU). (E) The SseA:SufE_Mtb_ AF3 model. (F) The SseA AF3 model. In (E) and (F) left panels shows the 0 replicate of the 5 produced AF3 models and the right ones the overlapping of the 5 models. N- and C-terminal domains of SseA are respectively colored in lavender and green, their connecting loop is in magenta and the Loop 185-201 is colored in orange. Side-chains of Cys245 and Ser251 at the active site are shown in sticks with carbons in brown. SufE_Mtb_ is shown in teal the loop containing the redox active Cys is highlighted in dark blue. Sidechains of key loops as well as those contributing to polar interactions are shown in sticks, while polar contacts are indicated by yellow (C) and grey (D-F) dashed lines.

Notably, as above indicated, the overall conformation of the 185-201 loop of SseA also shows major differences when comparing AF models with the crystal structure (**Fig. 10D-F**). In the crystal structure several interactions keep it in contact with the protein core: of note, Glu42 and Lys99 (at the N-terminal domain) hydrogen bond to the hydroxyl of Tyr193, while Arg246 (at the C-terminal domain) stablishes a cation-π interaction with His189 and forms a water bridge network that connects it to Glu42 through Glu249 (**Fig. 10D**). On AF models most of the mentioned residues at the protein core appear to maintain a very fix position, despite being slightly displaced from that of the crystal structure. However, the Arg246 side chain, in both the free and the complex AF3 models, shows a different conformation in each of the 5 produced models. The lack of a fix position for the Arg246 side chain relates with the loop twisting its two moieties regarding the protein core. Thus, Arg246 does not stack anymore to His189 that now points towards the protein surface, but makes a salt bridge with Glu196 (**Fig. 10E-F**). To be highlighted is the fact this Arg246 is just the following residue to the catalytic Cys245. On the contrary, Arg250, the residue preceding Ser251 assisting catalysis, shows a very fixed position in all evaluated structured by forming a salt bridge with Glu182 sit at a short helix preceding the 185-201 loop. Moreover, interaction of Tyr193 with the C-terminal domain of SseA become weaker in the AF3 free SseA model and fully disappear with SufE_Mtb_ is bound (**Fig. 10D-F**). Therefore, it might be envisaged that displacement of the water molecules at the entrance of the active site cavity and formation of the sulfur adduct at Cys245 of SseA may contribute to trigger displacement of Arg246. On its side, enhanced flexibility at the side chain of Arg246 would favor the unlocking of the mentioned 185-201 loop, and as a consequence SufE_Mtb_ complexation.

## Discussion

Tuberculosis (TB) is a critical mycobacterial infection that affects a wide range of mammals including humans. The main challenges in managing TB are the bacteria ability to enter a dormant, non-replicative state that resists drugs and immune response, along with the emergence of MDR strains, which have contributed to the recent rise in TB cases. There is a great need to discover new essential functions, in pathways different from those targeted by conventional antibiotics, to combat this disease.

In this frame we characterized SseA, a putative sulfurtransferase, overexpressed in MDR and XDR strains that has been demonstrated to participate in macrophage infection where it may play an essential role in oxidative stress resistance. By bioinformatic analysis, we discovered that SseA encoding gene presents a neighboring gene, Rv3284. The genomic proximity of this gene, which encodes for a SufE-like protein, suggested a potential functional relationship, prompting us to investigate Rv3284, herein further referred to as SufE_Mtb_.

Alignment of SufE_Mtb_ sequence with *E. coli* SufE revealed 60% sequence identity, including the conserved functional residue, Cys55, suggesting functional similarity between the two proteins. Surprisingly, this protein does not seem to be involved in Fe-S cluster biogenesis as its *E. coli* homologue, but in detoxification of cyanide [33].

To verify this latter hypothesis on SseA role in cyanide detoxification and SufE_Mtb_ ability to regulate it, we purified both SseA and SufE_Mtb_ as His-tagged fusion proteins that allowed for effective isolation. We obtained both proteins in a soluble monomer state and with high degree of purity.

We proved that SseA is a sulfurtransferase able to transfer sulfur from thiosulfate to cyanide and it shows it maximum activity at 37 °C. Beyond this temperature we observed a progressive decline in activity mostly mirrored by thermal instability and denaturation of the enzyme.

Having established the role of Mtb SseA, we proceeded to assess the role of SufE_Mtb_ considering their gene neighborhood, a reliable indicator of protein working together in a specific metabolic pathway. We proved indeed that SufE_Mtb_ promotes a considerable increase in the sulfur transfer activity of SseA. It is important to highlight this is the first time such ability is observed for a SufE-like homologue, whereas it is well characterized in the Suf operon where SufE is essential to promote SufS activity to carry out Fe-S cluster biogenesis[34].

Moreover, not all the enzymes of the sulfurtransferase family require an activator and, indeed, this characteristic seems unique to *Mycobacteria*.

To evaluate and further characterize whether the ability of SufE_Mtb_ to activate Mtb SseA resulted from direct protein-protein interaction, we applied different biophysical approaches that revealed the direct interaction with a moderate affinity (low µM range) and a 1:1 binding stoichiometry. It is worth noting that existing literature has already highlighted the functional differences between these domains in terms of sulfotransferase activity [35]. Typically, only the C-terminal domain presents the catalytic Cys residue necessary for enzymatic function. Multiple sequence alignment analysis reveals that SseA presents a clear sequence homology that is maintained across the represented domains, but certain sequences lack a disordered loop in the C-terminal domains. Indeed, many representatives of *Mycobacteria* present this extra loop (aa 185-201) that is absent, for example, in *E. coli*. This loop is present mainly in *Mycobacteria* where most likely the sulfurtransferase SseA requires to be activated by SufE-like protein.

We generated models of the SseA:SufE_Mtb_ interaction using AF2 and AF3 to investigate the structural basis of their interaction and the potential impact on SseA activation at the molecular level. The two models were virtually identical, suggesting a high level of reliability. Both models indicate that SufE-like proteins primarily interact with the non-catalytic N-terminal domain of SseA. The only contact with the catalytic C-terminal domain is mediated by a specific loop (residues 185–201).

These structural features could explain the experimentally observed 1:1 stoichiometry between SseA and SufE_Mtb_ proteins, as the extended loop may sterically or functionally constrain additional interactions.

Moreover, the 185-201 loop in the C-terminal domain and the region where SufE_Mtb_ binds in the N-terminal domain both appear to hinder access to the active site containing the SseA’s reactive cysteine. Upon SufE_Mtb_ binding, a conformational change has to occur, allowing access to the active site. Notably, both the 185-201 loop and SufE_Mtb_ catalytic cysteine region exhibit low-confidence structural predictions and increased mobility within the complex, a surprising observation, as protein complex formation generally stabilizes each binding partner. These insights suggest that the interaction induces a dynamic structural shift that enables SseA’s catalytic function.

Considering the very low prediction confidence of the SseA 185-201 loop within the SseA:SufE_Mtb_ complex, we suppose that its position can highly vary to allow Cys55 of SufE_Mtb_ to approach the SseA active site (**Fig. 9C**). In fact, in SufE_Mtb_, the loop containing Cys55 and the portion of the helix in contact with SseA also show less confidence than the rest of the structure. This further agrees with the SufE_Mtb_ loop allowing flipping of Cys55 from buried towards solvent exposed conformations to become accessible to the SseA active site. In addition, it might be envisaged that rearrangement of the contacts at the entrance of the active site cavity of SseA upon formation of the sulfur adduct, may be a factor contributing to trigger displacement of the mentioned 185-201 loop by Arg246 destabilizing its locking from the protein core (**Fig. 10D-F**). Thus, the presented models envisages a potential early activation stage through the SseA:SufE_Mtb_ interaction, which brings the SufE_Mtb_ loop in proximity with the catalytic site of SseA and prepares for subsequent catalysis.

## Materials and methods

### Protein Production

The synthetic gene of SseA from Mtb was ordered from IDT, Inc. (Coralville, IA, USA), as codon optimized sequence for expression in *E. coli* cells and cloned into the pET-based vector pETM-20 [17], using NcoI and NotI restriction sites, as a His-tagged thioredoxin (TrxA) fusion protein comprising a tobacco etch virus (TEV) protease cleavage site.

Mtb SufE-like protein (hereinafter identified as SufE_Mtb_) was amplified by PCR from a pET-DUET plasmid encoding for the protein already available in the laboratory. The PCR products were checked by 1% agarose gel electrophoresis and purified by a QIAquick-spin PCR purification kit (Qiagen, Hilden, Germany), double digested by NcoI and NotI, and ligated into a pET-24 expression vector placing SufE under lactose control and in frame with a His-tag, glutathione S-transferase (GST) and TEV protease site at the 5’ end. The cloned sequence used in this study was controlled by DNA analysis (BMR Genomics, Padua, Italy).

*E. coli* BL21-CodonPlus (DE3)-RP competent cells were transformed using the supplier’s protocol (Agilent Technologies, Ketsch, Germany). 100 µl of cells and 2.0 μl of a 1:10 dilution of the β-mercaptoethanol provided with the cells were incubated on ice for 10 min. DNA (10 ng) was added and cells were incubated for 30 min on ice, heat-shocked at 42 °C for 20 s and cooled on ice for 2 minutes. 900 µL of preheated (37 °C) SOC medium were added to the transformation reaction and incubated at 37 °C for 1 hour under gentle shaking. 200 µL of cell suspension were plated on selective media and incubated overnight at 37 °C

A pre-inoculum was prepared by picking a single colony from a plate, that was inoculated into 20 ml of 2xYT medium supplemented with the appropriate antibiotic (100 mg/L ampicillin for SseA, and 30 mg/L kanamycin for SufE_Mtb_). The culture was incubated overnight at 37°C with shaking at 200 rpm.

SseA and SufE overexpression were obtained by pouring the pre-inoculum into 2xYT medium supplemented with the appropriate antibiotic, and grown at 37 °C until an optical density (OD) at 600 nm of 0.6-0.8 was reached, followed by induction for 3-4 h by the addition of 0.5 mM isopropyl β-d-thiogalacto-pyranoside (IPTG) at 24 °C. The cell pellet was harvested and frozen. Frozen cells were thawed in a lysis buffer (20 mM Tris-HCl buffer, pH 8.0, 150 mM NaCl, 10 mM 2-mercaptoethanol, 30 mM imidazole, 5% (v/v) glycerol, 10 mg/L lysozyme, 1 mM phenylmethylsulfonyl fluoride (PMSF), 4.2 U/mL DNase) and subsequently sonicated and centrifuged. The proteins were purified by Ni-NTA affinity chromatography and eluted with 20 mM Tris-HCl buffer, pH 8.0, in the presence of 150 mM NaCl, 10 mM 2-mercaptoethanol, 300 mM imidazole. Purified proteins were cleaved from the tag by TEV protease and dialyzed overnight at 4 °C against 20 mM Tris-HCl buffer, pH 8.0, in the presence of 150 mM NaCl, and 10 mM 2-mercaptoethanol. Final purification was achieved by a second step of Ni-NTA affinity chromatography. When necessary, further purification was carried out by gel filtration chromatography on a ENrich SEC650 column (BioRad, Milan, Italy). The purity of the recombinant protein was checked by SDS-PAGE after each step of the purification.

Purified proteins were quantified at 280 nm (SseA ε_280_ 71515 M^−1^ cm^−1^, SufE ε_280_ 1490 M^−1^ cm^−1^) and flash-frozen.

### SDS-PAGE

Protein samples were diluted at a 1:1 ratio with denaturing buffer (0.125 M Tris-HCl, pH 6.8, 50% glycerol (v/v), 17 g/L SDS, 0.1 g/L bromophenol blue) in the presence of 2-mercaptoethanol and heated at 100 °C for 5 min. Electrophoretic runs were carried out in a MiniProtean Tetra Cell apparatus (Bio-Rad, Milan, Italy) on a 12% acrylamide-bisacrylamide (37.5:1 ratio) gel, using a Tris/glycine buffer system at pH 8.3. Gels were stained with GelCode™ Blue Safe Protein Stain (Thermo Fisher Scientific, Milan, Italy).

### Enzymatic assay

SseA thiosulfate sulfurtransferase activity was assessed by a discontinuous method that quantitates the thiocyanate produced by the enzymatic activity and is based on the absorption of the ferric thiocyanate complex at 460 nm (modified from Sorbo [18]).

Briefly, enzymatic activity measurements were performed at 37 °C in the presence of 10 µM SseA, 50 mM KCN, 1 mM DTT, and 50 mM KH_2_PO_4_, pH 8.6. Samples were incubated 30 minutes at 37 °C and the reaction was started by adding 50 mM Na_2_S_2_O_3_. To test the effect of SufE on SseA enzymatic activity, different concentrations of SufE (0, 10, 25, 50, 100 µM) were added during the 30 minutes incubation time before adding the substrate.

The assays lasted 30 min, and one unit of enzyme was defined as the amount of enzyme that produces 1 µmol thiocyanate per minute. At specific time points, enzymatic reactions were stopped by adding formaldehyde at a final 3.7% concentration, followed by the addition of 50 µL of 0.245 M Fe(NO_3_)_3_ ‧ 9 H_2_O (in 15% HNO_3_) that led to the formation of ferric thiocyanate. The reaction was incubated for 10 minutes and centrifuged at 14000 rpm, and 200 µL of supernatant was analyzed in a microplate absorbance reader (EnSight, Perkin Elmer, Milan, Italy).

Enzymatic activities at different temperatures were performed incubating and performing the assay at the reported temperature.

To determine the optimal temperature for activity, measurements were performed at the following temperatures: 4 °C, 16 °C, 25 °C, 37 °C, and 45 °C. This comprehensive temperature range allowed the identification of the temperature at which the highest level of activity was observed, providing insights into the performance of the system under various conditions. Each measurement was carried out at least in triplicate.

### SseA thermal stability

The thermal unfolding analysis was performed by using a Tycho instrument (NanoTemper Technologies, Munich, Germany). Measurements were carried out in the presence of 20 µM protein, using Tycho standard capillaries. Data were analyzed with the MO Affinity Analysis Software v2.1.3 (NanoTemper Technologies, Munich, Germany).

### MST measurements

Binding affinity between Mtb SseA and SufE_Mtb_ was investigated by MST using the Monolith NT.115 instrument (NanoTemper Technologies, Munich, Germany), using the Nano red LED at 100% excitation power and the MST power at 40%. About 0.1 nmol of SseA were labeled using the RED-NHS labeling second generation kit (MO-L011, NanoTemper Technologies, Munich, Germany) following the procedure recommended by the manufacturer. The dye carries a reactive NHS-ester group that covalently binds to primary amines (lysine residues). Labeled SseA concentration was ∼3 μM, with a degree of labeling of ∼0.5. Binding measurements were carried out at 25 °C in 50 mM sodium phosphate buffer, pH 7.8, containing 0.05% Tween-20, in the presence of 10 nM SseA and increasing concentrations of SufE_Mtb_ (from 1.5 nM to 50 μM). Fitting curves were obtained by the MO Affinity Analysis v3.0.5 software (NanoTemper Technologies, Munich, Germany).

### Isothermal titration calorimetry measurements

The calorimetric titrations were performed in a high-sensitivity automated Auto-iTC200 microcalorimeter (MicroCal, Malvern-Panalytical, Malvern, UK). A 10 µM SseA solution in the calorimetric cell was titrated with 100 µM SufE solution. Previously, the two protein solutions were chemically matched through fresh buffer exchange with PBS, pH 8.0 (phosphate 20 mM, NaCl 150 mM, DTT 2 mM). A series of 2-μL injections were performed with spacing of 150 s, stirring speed of 750 rpm, and reference power of 10 μcal/s. The heat evolved after each ligand injection was obtained from the integral of the calorimetric signal and normalized by the moles of protein injected. The heat due to the binding reaction between the inhibitor and the enzyme was estimated as the difference between the observed heat of reaction and the corresponding heat of dilution by including an adjustable term in the fitting routine accounting for the background injection heat. Applying a model considering a single binding site, the interaction parameters were estimated through non-linear least-squares fitting analysis: *K*_a_ (*K*_d_ = 1/*K*_a_) association constant; Δ*H*, interaction enthalpy; and *n*, apparent stoichiometry.

### Bioinformatics analyses

Amino acid sequences of known and orthologues rhodanese and rhodanese-like proteins were aligned using the ClustalW method [19]. The selected UniProt IDs with the corresponding species and protein name are described in Table 1.

**Table 1-.** Protein name, species and UniProt IDs in Fig. 7.

| Name | Species | UniProt ID |
| --- | --- | --- |
| Mtu_SseA | Mycobacterium tuberculosis | P9WHF6 |
| *Mle_THT3* | *Mycobacterium leprae* | Q50036 |
| *Mth_TST* | *Mycobacterium thermoresistibile* | E2QRA0 |
| *Mtu_CysA2* | *Mycobacterium tuberculosis* | P9WHF9 |
| *Mle_THTR* | *Mycobacterium leprae* | Q50036 |
| *Ser_THTR* | *Saccharopolyspora erythraea* | P16385 |
| *Avi_RhdA* | *Azotobacter vinelandii* | P52197 |
| *Eco_MST* | *Escherichia coli* | P31142 |
| *Mtu_THT3* | *Mycobacterium tuberculosis* | P9WHF4 |
| *Mth_THTR* | *Methanothermobacter thermautotrophicus* | O26719 |
| *Eco_YnjE* | *Escherichia coli* | P78067 |

### Building Ssea:SufE interacting models by AlphaFold

Structural models for the Mtb SseA:SufE interaction were produced by using the AlphaFold Multimer Colab v2.3.2 (AF2) (activating the Run_relax and Relax_use_gpu options and with a Multimer-max-number of cycles of 20) as well as the AlphaFold 3 (AF3) server [20–22], by using the corresponding protein sequences (UniProt ID P9WHF7 and P9WGC3 respectively). For comparison of potential structural changes regarding the free proteins, the coordinates of free SseA were taken from both the crystallographic structure with PDB ID 3HZU and from the AlphaFold Database (AFDB) with ID AF-P9WHF7-F1, while the structural model for SufE_Mtb_ was taken from the AFDB with ID AF-P9WGC3-F1 [23]. In addition, SseA and SufE_Mtb_ models were also constructed, for comparison by using the AF3 server. Models were analyzed using PyMOL [24].

## Author contributions

S.A. conceived and supervised the project. A.F. performed and analyzed the MST and thermal stability measurements. A.F. and G.D.N. wrote the original draft of the manuscript. G.D.N. and E.R. purified the proteins and investigated the enzymatic kinetics. E.S. performed bioinformatic analyses. A.V.C. performed and analyzed the ITC measurements. A.R. and R.B. performed the cloning and expression of the proteins. M.Me. performed the computational studies of the interaction between the proteins. S.O.B. and M.Ma helped with the experimental design, contributed to the interpretation of results and to the final version of the manuscript. All authors provided critical feedback and helped in shaping the research and the manuscript.

## Acknowledgements

This work has been funded by the University of Turin (Ricerca Locale Grants ADIS_RILO_21_01, OLIS_RILO_21_02, MARM_RILO_22_05; Grant for Internationalization MARM_GFI_22_01_F_01); the Spanish State Research Agency and FEDER (MCIN/AEI-FEDER, Grant PID2022-136369NB-I00); and the Government of Aragón-FEDER (Grant E35_23R). This work received funding in the frame of: Project INF-ACT “One Health Basic and Translational Research Actions addressing Unmet Needs on Emerging Infectious Diseases PE00000007”, PNRR Mission 4, EU “NextGenerationEU”-D.D. MUR Prot.n. 0001554 of 11/10/2022, CUP B53C20040570005; and Project PRIN 2022 PNRR, funded by the European Union, “NextGenerationEU”, Mission 4 Component 2. CUP B53D23033100001, Prot. P2022JE8FN-“FIGHT_TB:FIndinG High-grade anTigens towards an innovative TB vaccines” - D.D. MUR Prot. n. 0001363 of 1 September 2023. This work was also supported by MOSBRI, a project receiving funding from the European Union Horizon 2020 Research and Innovation Programme under grant agreement No. 101004806 (MOSBRI-2022-134 and MOSBRI-23-185).

## Notes

**Conflicts of interest**: The Authors declare no competing interests.

### Competing Interest Statement

The authors have declared no competing interest.

